# GRID-seq assisted prediction of transcription factor binding motifs

**DOI:** 10.1101/429332

**Authors:** Werner Pieter Veldsman

## Abstract

Experimental validation of computationally predicted transcription factor binding motifs is desirable. Increased RNA levels in the vicinity of predicted protein-chromosomal binding motifs intuitively suggest regulatory activity. With this intuition in mind, the approach presented here juxtaposes publicly available experimentally derived GRID-seq data with binding motif predictions computationally determined by deep learning models. The aim is to demonstrate the feasibility of using RNA-sequencing data to improve binding motif prediction accuracy. Publicly available GRID-seq scores and computed DeepBind scores could be aggregated by chromosomal region and anomalies within the aggregated data could be detected using mahalanobis distance analysis. A mantel’s test of matrices containing pairwise hamming distances showed significant differences between 1) randomly ranked sequences, 2) sequences ranked by non-GRID-seq assisted scores, and 3) sequences ranked by GRID-seq assisted scores. Plots of mahalanobis ranked binding motifs revealed an inversely proportional relationship between GRID-seq scores and DeepBind scores. Data points with high DeepBind scores but low GRID-seq scores had no DNAse hypersensitivity clusters annotated to their respective sequences. However, DNase hypersensitivity was observed for high scoring DeepBind motifs with moderate GRID-seq scores. Binding motifs of interest were recognized by their deviance from the inversely proportional tendency, and the underlying context sequences of these predicted motifs were on occasion associated with DNAse hypersensitivity unlike the most highly ranked motif scores when DeepBind was used in isolation. This article presents a novel combinatory approach to predict functional protein-chromosomal binding motifs. The two underlying methods are based on recent developments in the fields of RNA sequencing and deep learning, respectively. They are shown to be suited for synergistic use, which broadens the scope of their respective applications.

## Introduction

To consider the effect that the presence of an increased concentration of RNA in the vicinity of a promoter may have on a protein binding to a promoter, an approach that combines an estimate of RNA affinity and transcription factor (TF) affinity for the same given sequence is required. Two recent papers have each presented a new approach that have the potential to be applied towards this goal in combination. Firstly, there is the global RNA interactions with DNA by deep sequencing (GRID-seq) approach(1), which allows a user to quantify genome wide binding of RNA to chromatin in a manner that results in data similar to what might be obtained from model-based analysis of ChIP-Seq (MACS)(2). In other words, a BED file is generated that contains chromosomal regions of an arbitrary length of 1000 nucleotides, annotated with a score that indicates the extent to which RNAs bind to a given sequence. The second approach, called DeepBind(3), uses a deep learning approach to discover TF binding motifs in a given sequence. The models with which DeepBind’s binding motif predictions are made were trained using either ChIP-seq, SELEX or RNA-binding protein (RBP) experiments. This is a defining characteristic of DeepBind that diversifies the provenance of the data that its models are derived from.

Although DeepBind and GRID-seq are new experimental techniques, the principals on which they are based have already been validated by a significant amount of research. As an example, the DeepBind method employs a specific type of deep learning neural network called a convolution neural networks (CNNs). CNNs have been used to predict RNA secondary structure(4) and impute DNA methylation(5) to name but a few. GRID-seq for its part has the major advantage over previous methods in that it can be used to detect nascent RNA-chromatin interaction on a genome wide scale. Analyzing nascent RNA is preferable over analyzing total cellular RNA since it is believe to be a less biased approach that provides the researchers with better resolution over cellular variables such as RNA stability and transcription rate(6).

The aim of combining DeepBind and GRID-seq is to derive results that may assist researchers in study target selection. The desired outcome is achieved by simultaneously considering the influence that scores from of each of the respective methods (GRID-seq and DeepBind) has as a single distance function based on the distribution of their scores, and then ranking sequences that contain predicted binding motifs according to this new combined score. The intention is to predict protein-chromatin binding motifs with an indication as to whether the predicted motifs might be bound by proteins in an environment with heightened levels of RNA, which could in turn indicate whether the binding motif is functional.

## Methods

### Sourcing GRID-seq data, DeepBind models, and related sequences

Candidate transcription factors were found in GRID-seq supplementary data by manually filtering gene names published by Li *et al*. (1) The criteria for selecting a gene name from the GRID-seq data was that a DeepBind model matching a gene name was available at http://tools.genes.toronto.edu/deepbind/ Multiple matches were found using this approach. An arbitrary decision was made to select *CUX1* related data for the purposes of illustrating the method discussed in this paper. The first step after confirming that both GRID-seq data and a DeepBind model existed for a gene of interest, was to retrieve GRID-seq genomic intervals and accompanying RNA-binding scores from http://fugenome.ucsd.edu/gridseq/ This data was found by matching the gene identifier to file names in the MM.1S dataset. GRID-seq results for *CUX1* were extracted from file gridseq. MM1S.ENSG00000257923.bdg The actual DNA sequences were downloaded by concatenating these ranges to an http get request directed at http://togows.org/api/ucsc/hg38/

### Predicting protein-chromatin binding motifs using a DeepBind model

The sequences obtained in the previous step each had a length of 1000 nucleotides. These sequences had to be cut into 20 shorter sequences of length 50 to comply with DeepBind specifications. DeepBind was downloaded from http://tools.genes.toronto.edu/deepbind/ The shortened sequences and the CUX1 model identifier (D00329.002) were passed as input to the DeepBind modeler. A file by the name README.txt file is bundled with the DeepBind modeler and contains usage guidelines and handy examples. To keep scoring regimes comparable, the maximum DeepBind score in each bin of 20 subsequences were used as the matching DeepBind score for each of the GRID-seq scores associated with a 1000 base pair genomic region.

### Score integration and sequence ranking

GRID-seq and DeepBind scores per genomic interval were recoded into single variables using the multivariate mahalanobis distance method of the R package anomalyDetection(14). GRID-seq and DeepBind scores together with their respective means and co-variances were passed to *anomalyDetection’s* mahalanobis function whilst leaving all other parameters at their default values. To improve on the resolution of data points, an arbitrary decision was made to keep 1% of the highest ranking Mahalanobis scores (60 out of 6069) for further analysis. A hamming distance matrix was created from sequences associated with the top 60 mahalanobis scores using the R package *DECIPHER(15)*. Similar matrices were created for the top 60 sequences ranked by DeepBind scores and 60 randomly selected sequences. These three matrices were then tested for correlation in turn using a Mantel test implementation bundled with the R package *ade4*(16) with the default permutation value of *ade4’s* mantel.rtest method changed to 10,000. The null hypothesis for each paired test of the three matrices were that there is no relationship between any two given matrices at α = 0.05 Data parsing was carried out using the R packages *data.table*(17) and *reshape2*(18). In all cases scores were visualized using the R package *ggplot2*(19), and where applicable, *gridExtra*(20).

### Test for robustness

In the main example discussed in this paper (Fig 1), CUX1 transcription factor binding motifs were predicted using a DeepBind model that was trained on SELEX data. To test the robustness of GRID-seq assisted protein-chromatin binding motif prediction as well as the approach’s tolerance to changes in DeepBind model architecture, two further transcription factors were analyzed. They were TBL1XR1 and MTR3. DeepBind models for these two TF’s were constructed using ChIP-seq and RBP assay data, respectively. Scores derived using each of the three DeepBind models were visually compared to each of the GRID-seq associated DNA sequences. The results of these further tests are illustrated in Fig 2.

**Fig 1:**
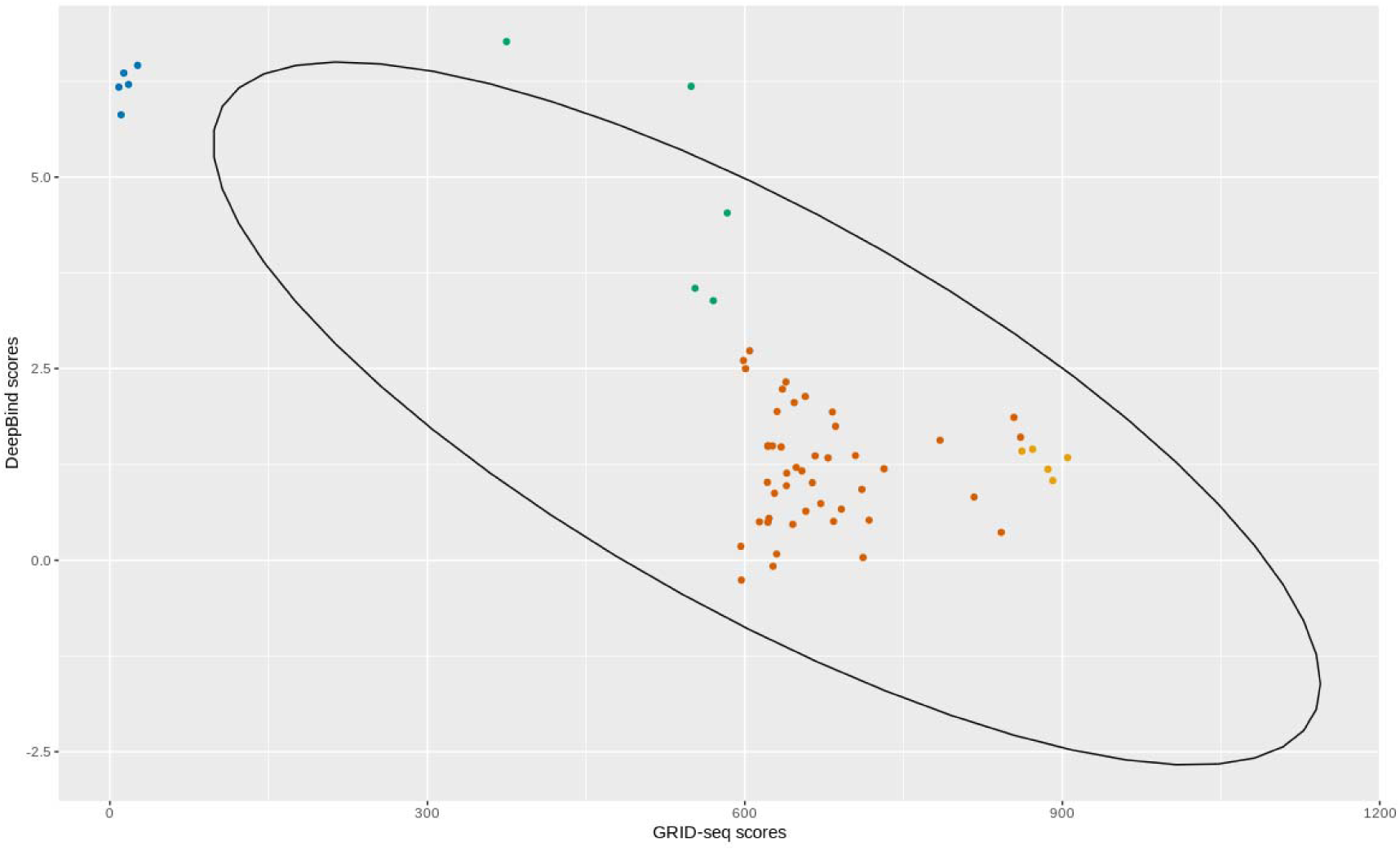
Mahalanobis ranked binding motifs. Two predicted binding motif scores in the green cluster, directly above and outside the confidence ellipse, are noticeably separated from both high DeepBind scores (blue cluster) and high GRID-seq scores (orange and yellow clusters). These two data points are separated from other high DeepBind scores in the blue cluster because moderate levels of RNA have been detected in the vicinity of their associated binding motifs.

**Fig 2:**
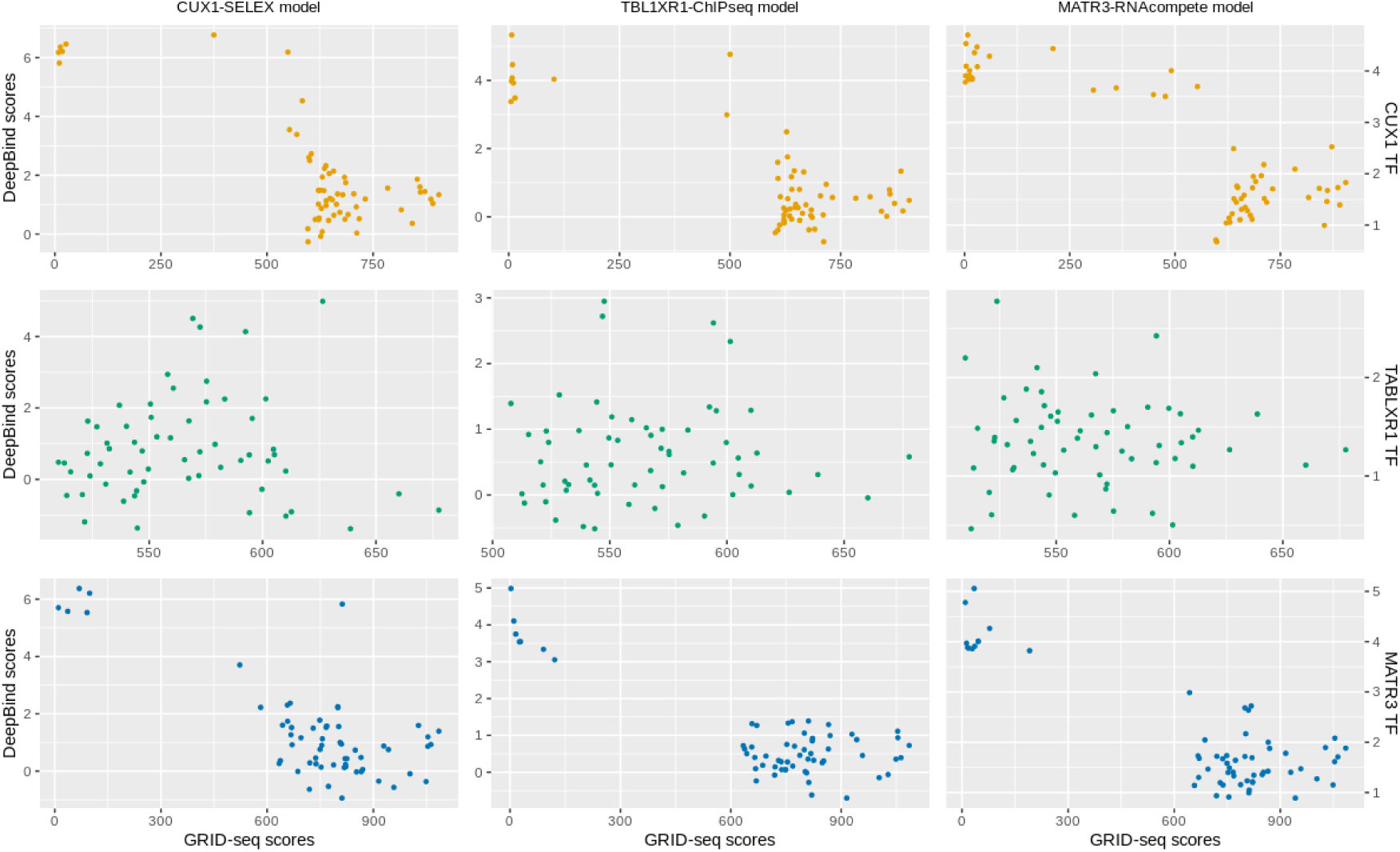
Permuted test of robustness using three models and three transcription factors. GRID-seq data plotted against DeepBind data for three transcription factors and three models derived from different source data. There seems to be a general trend that the higher a DeepBind score is, the lower the level of RNAs are in the vicinity of the motif predicted by DeepBind. However, it is the outliers that deviate from this pattern that are of interest due to their higher potential to represent functional binding motifs.

## Results

### Mahalanobis distance analysis reveals candidate binding motifs as outliers

GRID-seq scores in the top 60 mahalanobis ranking had a minimum value of 8.47 as compared to a minimum value of 2.03 in the entire GRID-seq score population. Similarly, DeepBind scores in the top 60 mahalanobis ranking had a minimum of −0.26 as compared to a minimum value of −1.97 in the entire DeepBind score population. Maximum values for GRID-seq and DeepBind scores were 905.06 and 6.77 in the mahalanobis subsets. Maximas in the subsets were identical to those in the entire populations. The minima and maxima properties therefore indicated that the top 1% mahalanobis ranked scores were qualitatively representative of their respective populations. Plotting the top 1% mahalanobis ranked scores by their constituent GRID-seq and DeepBind scores illustrated how mahalanobis ranking can be applied to increase the resolution of realistic candidate binding motifs (Fig 1)

### RNA levels are associated with significantly dissimilar sequence ranking

A statistical test to determine whether the order of DNA sequences (and therefore the order of predicted motifs) were significantly affected by choice of scoring method, revealed that Mahalanobis ranked sequences were significantly different to DeepBind ranked sequences (p=0.45). Mahalanobis ranked sequences were also significantly different to randomly ranked sequences (p=0.75) and DeepBind ranked sequences were significantly different to randomly ranked sequences (p=0.80). These results statistically confirm that the ranking of DNA sequences (that contain predicted binding motifs) are significantly changed by the choice of scoring method. The results are as expected; however, it validates the motif prediction power of DeepBind for this specific case in that the sequences that are hypothesized to contain binding motifs are statistically proven to be compositionally different to random sequences. In addition, the lower p-value with which DeepBind-ranked and Mahalanobis-ranked sequences were deemed to be statistically different from each other suggests that the underlying sequence composition that leads to the prediction of a binding motif is not disproportionally affected by either GRID-seq data or DeepBind data. If it had been disproportionally affected, the p-value would have either shown an insignificant difference (p < 0.05) or would have been similar to the p-values obtained by tests on randomly ranked sequences.

### GRID-seq and DeepBind scores have an inversely proportional property

A comparison of GRID-seq assisted transcription factor motif prediction scores for model types against different sequence sets show that clustering of high DeepBind scores and GRID-seq scores are dependent on the strength of the respective scores (Fig 2). From these nine graphs, it seems in six out of the nine cases that the higher the average DeepBind scores are, the higher the average GRID-seq scores are. However, when looking at the individual data points, it becomes apparent that there is a tendency that the higher an individual GRID-seq score is, the lower its accompanying DeepBind score is. In other words, an inversely proportional relationship between GRID-seq scores and DeepBind scores seems to become more pronounced as their respective values increase.

## Discussion

Discovering transcription factor binding motifs *in silico* requires follow up work to establish whether binding takes place *in situ*. This is the case because not all potential binding sites are bound by transcription factors in the cellular environment. These motifs that are compositionally valid but never bound are the false positives of computational motif prediction. It has been suggested as early as 2002 that expression data can be used to validate functional binding motifs(7). Employing GRID-seq data would therefore be a realization of using such an approach. Many software solutions have already been developed for computational prediction of binding motifs. A 2006 paper by Tompa et al. evaluated the performance of thirteen of these tools(8). In Tompa’s discussion, emphasize is placed on using complementary tools for motif prediction and to investigate a “top few” predicted motifs instead of just the most significantly predicted motif.

But the question as to how RNA levels may contribute to confirming a binding motif’s functional role has multiple answers. A protein’s function can be changed when bound by RNA. As a result, a transcription factor bound by RNA may promote, inhibit or completely uplift transcription. Transcription factors (TFs) have the ability to bind to RNA as well as to DNA. Binding of RNA to a nuclear protein can override the protein’s localization signal, leading to its export from the nucleus(9). It has also been shown that a well-known TF, p53, has two distinct nucleotide binding domains: one with high specificity for motifs in the promoter region of genes targeted by p53, and another domain that exhibits non-specific binding to, amongst others, single stranded nucleotide sequences. TFIIIa is an example of a typical zinc-fingered TF that binds both DNA and RNA(10). Given then that RNA has been demonstrated to have the ability to interfere with the localization signal of a protein and to have the ability to bind to transcriptional initiation factors in a non-specific manner, it is reasonable to assume that there exists a large set of RNAs in the nucleus that can directly affect gene expression that is in turn under the control of a TF. More specifically, the binding of a transcription factor to a promoter region could therefore be under the regulation of an unknown number of RNAs. And since the binding may be non-specific, it follows that if the concentration of RNA in the vicinity of a promoter is high, there must be a higher probability that the binding of the TF could be regulated. If not by the possibility that the given TF has non-specific RNA binding affinity, or that increased RNA levels indicate 3d proximity of some sequentially distant enhancer/repressor, then at least by the increased likelihood of physical obstruction arising from the quantity of RNA molecules in the vicinity of the promoter. This rationale forms the basis of the method that is presented in this paper. In this sense, the method purposed here does not draw specific conclusions about the nature of the association between RNA and predicted protein binding motifs but acts as a mechanism by which candidate binding motifs can be selected for further study.

When switching around cause and effect, the proximity of RNA to a predicted binding motif could instead be taken as an indication that transcription is taking place due to regulatory activation of a downstream gene. In that case, the flow of information as described by the central dogma(11) suggests that it is the presence rather than the mechanism of action of RNA that gives a clue as to whether a binding motif is functional. The inverse proportional relationship of GRID-seq scores and DeepBind scores that becomes apparent when viewing Fig 2 seems to indicate that the more likely it is that a given sequence contains a binding motif, the less RNA are bound to the chromatin in its vicinity. This does not support or reject the central dogma notion of confirming binding motif validity by the presence of RNA, because although most of the motifs predicted with high significance exhibit the inversely proportional relationship, a few high significance motifs do not. It does however allow for a novel categorization of the “top few” significant motifs – i.e. those that are associated with heightened levels of RNA and those that are not.

The contribution that GRID-seq data can make when assessing the functionality of a predicted binding motif becomes apparent in Table 1. This table lists the top 5 predicted motifs ranked by their DeepBind scores (blue cluster on Fig 1), the top 5 ranked by their GRID-seq scores (yellow cluster on Fig 1), and five other predicted motifs (green cluster on Fig 1) that are discernable on Fig 1 as being separated from both the DeepBind and Grid-seq clusters. What the table shows is that even though DeepBind might predict certain motifs with high significance, when the DNAse I activity of these top 5 sequences ranked by DeepBind score is examined, there appears to be no DNAse I cluster annotated to them (when viewed using the UCSC Genome browser(12)). The DNAse cluster column in Table 1 is a binary field that either confirms that cluster boxes have been observed when viewing the genomic regions in the UCSC Genome browser or that cluster boxes have not been observed. DNAse hypersensitivity typically indicates that chromatin is in an accessible state and has been extensively used to identify regulatory elements including promoters(13).

**Table 1:** DNAse hypersensitivity characteristics of sequences containing predicted motifs.

|  |  |  |  |  |  |  |  |
| --- | --- | --- | --- | --- | --- | --- | --- |
|  | GRID-seq score | DeepBind score | Mahalanobis distance | Chromosome | Start | Stop | DNase clusters |
| GRID-seq | 905.056 | 1.338331 | 47.59 | chr7 | 101994000 | 101995000 | No |
|  | 890.99 | 1.038863 | 46.24 | chr7 | 101993000 | 101994000 | No |
|  | 886.461 | 1.186810 | 45.61 | chr7 | 101991000 | 101992000 | Yes |
|  | 871.941 | 1.448033 | 43.90 | chr7 | 101992000 | 101993000 | No |
|  | 861.87 | 1.423735 | 42.82 | chr7 | 101990000 | 101991000 | Yes |
| DeepBind | 26.255 | 6.457482 | 27.66 | chr7 | 100805000 | 100806000 | No |
|  | 13.0953 | 6.356945 | 26.77 | chr7 | 104937000 | 104938000 | No |
|  | 17.6181 | 6.207847 | 25.23 | chr7 | 103866000 | 103867000 | No |
|  | 8.47069 | 6.174184 | 25.01 | chr7 | 135598000 | 135599000 | No |
|  | 10.5831 | 5.812598 | 21.56 | chr7 | 93166000 | 93167000 | No |
| Combined | 583.454 | 4.531703 | 27.71 | chr7 | 101816000 | 101817000 | Yes |
|  | 570.183 | 3.386700 | 21.17 | chr7 | 101891000 | 101892000 | No |
|  | 552.997 | 3.548660 | 20.71 | chr7 | 101815000 | 101816000 | Yes |
|  | 549.316 | 6.183928 | 38.21 | chr7 | 101940000 | 101941000 | No |
|  | 374.84 | 6.767595 | 35.56 | chr7 | 101927000 | 101928000 | Yes |

Increased RNA levels in the vicinity of accessible chromatin makes intuitive sense if they are involved in regulating a functional binding motif or are transcribed in close proximity to a functional binding motif. In other words, since increased RNA levels are expected to be observed in the vicinity of a functioning gene (and by consequence it’s promoter and TF binding motifs), it could indicate that a given binding motif is more likely to be functional than a binding motif in a region that does not show increased RNA levels. It should be noted that if regulatory interfering RNAs were to be a significant constituent of an observed increased level of RNAs in a given chromosomal region, the assumption about the reason for their presence would be the opposite of the assumption for the cause of their presence under the central dogma.

## Conclusion

The approach presented here demonstrates that RNA-seq data can be used to substantiate binding motifs predicted by computational methods. It is shown that predicted binding motifs that have some RNA bound in their vicinity also on occasion have DNAse hypersensitivity annotated to their contextual sequences. This characteristic stood in contrast to the most highly ranked predicted binding motifs by computational prediction, which had no hypersensitivity annotated to their contextual sequences. DNAse hypersensitivity is widely accepted as indicative of transcriptional activity. The ability to distinguish between true positive and false positive binding motifs is therefore improved by this approach. In addition, this approach allows simultaneous consideration of both RNA-seq and DeepBind data using a novel statistical application of the mahalanobis distance function, which in turn allows binding motif candidates to be ranked based on the contribution of multiple variables. Future work could focus on experimenting with the inclusion of additional variables that may further improve the accuracy and specificity of predicted binding motifs.

